# Interspecific divergence of gene expression biophysics driven by both evolutionary systems drift and strong selective constraints on bursting rate

**DOI:** 10.1101/2025.11.24.690267

**Authors:** Catherine Felce, Joshua G. Schraiber, Madhumitha Krishnaswamy, Alexander L. Cope, Lior Pachter, Matt Pennell

## Abstract

Interspecific comparisons of cell-type-specific gene expression levels can provide information about the evolutionary processes that drove divergence between species. From these comparisons, it is now evident that the predominant mode of gene expression evolution has been stabilizing selection, both on the steady-state (mean) protein levels as well as on the mRNA levels with additional lineage-specific shifts resulting from directional selection. However, as all previous work has used bulk RNA measurements, it has been impossible to determine which of the many cellular processes that contribute to mean abundances are highly constrained and which are more evolutionary labile. Assessing this is further complicated by the expectation that components of complex systems will evolve over time independent of changes in the selective regime so long as the net output of a system (i.e., mean expression) remains near the evolutionary optima. This process, known as evolutionary systems drift (ESD), has been frequently invoked as a non-adaptive explanation for changes in cellular phenotypes but has never been quantitatively tested or accounted for in any statistical test for selective constraints. Here, we develop a new paradigm that addresses both of these open problems simultaneously. Using single-cell expression data and biophysical models, we estimate mRNA transcriptional bursting rates, splicing rates, and decay rates across multiple vertebrate species. We then derive new mathematical results that describe how these various biophysical parameters are expected to co-evolve under ESD and then test whether we need additional evolutionary constraints to explain the divergences in these parameters. We find evidence that the biophysical parameters are indeed evolving in a coordinated manner as predicted by ESD and that there are additional strong constraints on transcriptional bursting, likely as a consequence of selection to reduce noise in expression. More broadly, this work opens up a whole new approach for studying the evolutionary dynamics of complex cellular systems.

## Introduction

Gene expression divergence is understood to be a key determinant of phenotypic divergence between species [1–3]. Sequencing technologies enabled comparative analyses of gene expression across individuals and species, providing insights into the molecular and evolutionary mechanisms shaping gene expression evolution, with most studies focusing on mRNA expression evolution measured via RNA-seq. Numerous studies have investigated gene expression evolution across species, revealing both widespread stabilizing selection and lineage-specific adaptive shifts in steady-state (hereafter, mean) mRNA expression levels [4–8]. Recently, Cope et al. [9] used phylogenetic modeling to show that there is stabilizing selection on mRNA expression levels per se and not simply as a mechanistic consequence of stabilizing selection on protein levels [10]. Additionally, this data has been used to identify complex evolutionary patterns such as organ-specific evolution [6, 11], conserved gene regulatory modules [12], differential expression correlated with the emergence of complex phenotypes [13], and gene-by-gene coevolution of gene expression [14].

Despite the progress made in identifying patterns of gene expression evolution and the processes that drive them, these studies have been limited by their use of mean mRNA measurements. While mean expression levels are a useful measure for studying gene expression evolution, these obfuscate how expression levels evolve and which mechanisms of gene expression are most dynamic or most constrained by natural selection. Mean mRNA levels are determined by a large array of interacting cellular processes [15–20], which can make different contributions to between-species expression divergence.

To access information about these important regulatory mechanisms from transcriptomic data, we need new approaches that explicitly consider biophysical parameters. Principled biophysical modeling is crucial for extracting biological information from RNA-seq data [21]. The emergence of single-cell RNA-seq technologies has allowed for the estimation of key biophysical parameters, such as transcriptional burst size and frequency [22–24]. Single-cell snapshot experiments can effectively provide two ‘time points’, via the analysis of nascent and mature transcripts [25–27]. Such models have been used to estimate the relationship of transcriptional bursting to cell-cycle stage [28], for principled cell-type clustering [29], and to disentangle biological and technical noise [30]. The quantification of biophysical parameters from single-cell data opens up new routes for investigating mRNA expression evolution at the levels of transcriptional and post-transcriptional dynamics. With the increased availability of cross-species single-cell datasets, comparative analyses of biophysical parameters could reveal new insights into the evolution of gene expression on macroevolutionary timescales. In this work, we introduce a new paradigm for investigating the evolutionary interplay of different regulatory components. Instead of bulk RNA measurements, we use single-cell data to understand the evolution of biophysical parameters across vertebrates (Figure 2).

In order to do this, we needed to confront another major roadblock faced by any evolutionary analysis of a complex system: change, not stasis, is the expectation. Even if the selective regime remains constant, the underlying components of a complex system can evolve so long as the net output (here, mean expression level) remains constant. This idea, now broadly referred to as “evolutionary systems drift” (ESD), has been invoked (though not always by name) as a non-adaptive explanation for much interspecific variation in complex cellular systems, such as enhancer sequences [31], gene regulatory networks [32, 33], gene expression levels [9, 10, 34], patterns of protein phosphorylation [35], and Transcription-Factor/Transcription-Factor-Binding-Site interactions [36]. And there are many additional cases where compensatory evolutionary changes in gene regulation have been observed, even if they have not been interpreted as instances of ESD [37–39]. For example, experimental work using a two-species yeast hybrid found that changes to mRNA degradation rates were often accompanied by opposite-effect changes in transcription rate [40].

But critically, ESD has been primarily a verbal model and, as such, it has not been possible to quantify the expected amount or pattern of change under this model nor compare it to alternatives — which is essential for evaluating claims of adaptation. In this paper, we extend recent theoretical results [10, 41] and derive quantitative expectations for the co-evolution of system components under ESD where only the net output is under selection, selection on the individual components, and a combination of these scenarios. We then fit these alternative scenarios to the biophysical parameters we estimated from multi-species single-cell data. We find strong evidence that the biophysical parameters are indeed evolving in a coordinated manner, as predicted by ESD, but that evolution of gene expression is further constrained by strong selection on transcriptional bursting.

## Results

### Biophysical parameters of gene expression vary across vertebrate species

For our basic biophysical model, we use bursty transcription with the following dynamics:

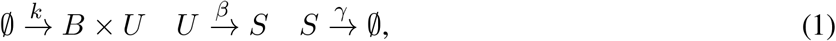

where transcriptional bursts [42] occur at a rate *k*, producing bursts of unspliced (*U*) transcripts with sizes distributed according to *B*, a geometric distribution with mean size *b* (a combined parameter of burst size and burst frequency). The unspliced transcripts are converted to spliced mRNA (*S*) at a rate *β*, and the spliced transcripts then decay at a rate *γ*. The ∅ symbol indicates that the transcript before transcription and after decay does not appear in the model.

We estimated per-gene values of the biophysical parameters *b, β*, and *γ* using Monod [43] (see Methods), an inference framework that maximizes the likelihood of single-cell spliced and unspliced RNA count data under this model. Applying Monod to spleen T-cell single-cell transcriptomics data from Jiao et al. [44] across the six retained vertebrate species yielded per-gene, per-species parameter estimates for 167 orthologous genes (**Fig. 1**).

**Fig 1:**
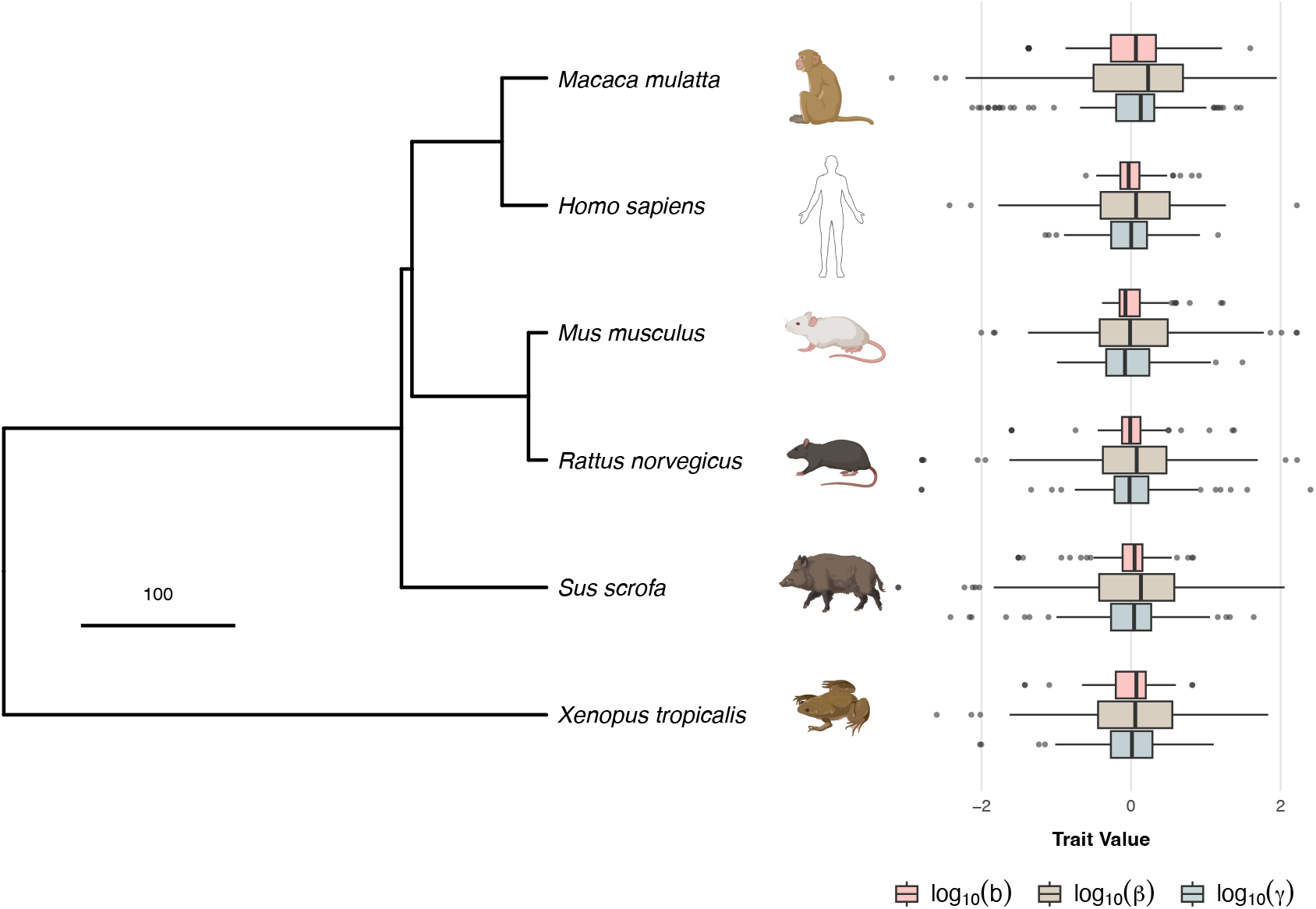
Biophysical parameters of transcription vary across spleen T cells from six vertebrate species. The phylogeny is taken from TimeTree [45] (scale bar in millions of years, Myr). Silhouettes taken from PhyloPic https://www.phylopic.org/. Boxplots at each tip show the distribution across 167 orthologous genes of the mean-centered log values of burst size *b* (pink), splicing rate *β* (grey), and mRNA decay rate *γ* (blue) estimated by Monod [43] from spleen T-cell single-cell RNA-seq data. Mean-centering reveals relative differences in biophysical parameters.

**Fig 2:**
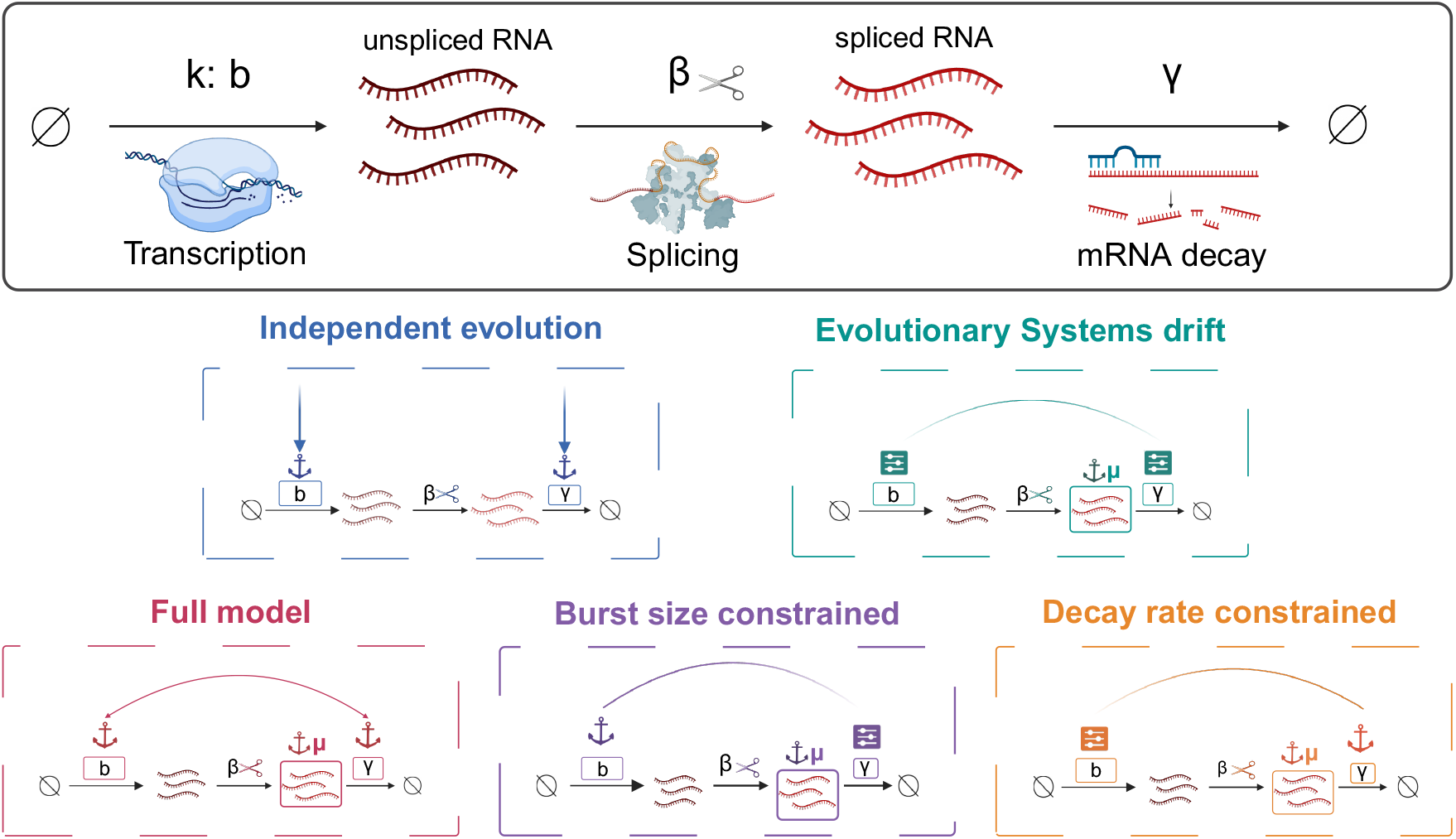
Competing evolutionary models for the joint evolution of transcriptional burst size and mRNA decay rate. The top panel shows the biophysical model of gene expression: transcriptional bursts (rate *k*, mean burst size *b*) produce unspliced RNA, which is spliced to mature mRNA at rate *β*, which then decays at rate *γ*. The five panels below depict the competing phylogenetic models tested in this study. In the Independent Evolution model, *b* and *γ* each evolve under stabilizing selection toward their own optima without coupling. In the Evolutionary Systems Drift (ESD) model, only mean expression log(*µ*) = log(*b*) − log(*γ*) is under selection, and *b* and *γ* co-evolve in a correlated but non-directional manner. The burst-size constrained, decay-rate constrained, and full models add direct stabilizing selection on one or both individual components on top of the ESD structure. Coloured anchors indicate stabilizing selection; the tuning box (implying flexibility) indicates a parameter that evolves freely. Figure created using Biorender

We focused our evolutionary analysis on burst size *b* and decay rate *γ* because together they determine the log mean spliced mRNA level,

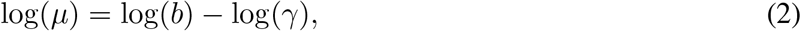

which is independent of the splicing rate *β* (with *γ* given in units of the burst initiation rate *k*); for notational simplicity, we denote log-transformed parameters by *ℓ*_*i*_, hereafter. Qualitatively, across most species we observe a wider range for the burst size *b* than for the decay rate *γ* **(Fig. 1)**, motivating our investigation into the selective forces governing their joint evolution.

### Evolutionary systems drift predicts coordinated but non-directional co-evolution of burst size and decay rate

A central theoretical contribution of this paper is to derive the expected joint evolutionary dynamics of *b* and *γ* under different hypotheses about which aspects of gene expression are under natural selection. We model *ℓ*_*b*_ and *ℓ*_*γ*_ as quantitative traits evolving along the phylogeny and derive that, under a broad class of stabilizing selection scenarios, their joint evolution follows a bivariate Ornstein-Uhlenbeck (OU) process (Eq. (4), Methods). This is the standard comparative biology model of stabilizing selection [46], in which traits are pulled toward an evolutionary optimum at a rate proportional to their displacement. The specific form of the selection matrix *H* in this model encodes which aspects of gene expression are under selection and whether the two parameters are evolutionarily coupled.

Starting from a general fitness function that allows stabilizing selection to act simultaneously on burst size, decay rate, and/or mean spliced expression (Eq. (3), Methods), and applying the classical quantitative genetics framework of Lande [47] to the bivariate system, we derive the forms of *H* for five biologically distinct selective scenarios. Extending prior theoretical work by Veller and Muralidhar [41] and Jiang et al. [10] on the marginal dynamics of individual system components, a key new result is the derivation that the joint evolution of *ℓ*_*b*_ and *ℓ*_*γ*_ under evolutionary systems drift (ESD) takes the form of a bivariate OU process with the singular selection matrix *H*_ESD_ (Eq. (5), Methods). Previous work had established that each component individually evolves as a Brownian motion (BM) under ESD, with no net directional pull on *ℓ*_*b*_ or *ℓ*_*γ*_ in isolation. We show that the joint process has a richer structure: its eigendecomposition (Eq. (6), Methods) reveals two orthogonal evolutionary modes. The first is a neutral axis along which *ℓ*_*b*_ and *ℓ*_*γ*_ drift in exact tandem at no fitness cost, corresponding to the marginal BM of previous work. The second is a constrained axis along which changes in *ℓ*_*µ*_ = *ℓ*_*b*_ − *ℓ*_*γ*_ are resisted by stabilizing selection, so that mean expression itself evolves as a scalar OU process (Eq. (7), Methods). The joint OU process captures both modes simultaneously, and crucially implies that changes in *ℓ*_*b*_ and *ℓ*_*γ*_ across species are not independent: a drift in burst size in one lineage generates a compensatory selective pressure on the decay rate. This correlated structure is the quantitative signature of ESD and is what makes it distinguishable from independent evolution in comparative data.

We derived four additional models representing alternative selective scenarios (Eqs. (8)–(11), Methods; **Fig. 2**). In the “independent” model (*H*_indep_), *ℓ*_*b*_ and *ℓ*_*γ*_ each evolve under stabilizing selection toward their own optima with no coupling between them. In the “burst-size constrained” model (*H*_*b*-constrained_), burst size is under direct stabilizing selection while decay rate is only indirectly constrained through its contribution to *ℓ*_*µ*_. In the “decay-rate constrained” model (*H*_*γ*-constrained_), the roles are reversed: decay rate is directly constrained and burst size adapts through ESD-type coupling. Finally, the “full” model (*H*_full_) combines direct selection on both *ℓ*_*b*_ and *ℓ*_*γ*_ with ESD-type coupling from selection on *µ*. All five models share a common mutational structure in which new mutations affect *ℓ*_*b*_ and *ℓ*_*γ*_ independently (Eq. (12), Methods).

Because we have data from only six species but estimates of biophysical parameters from 167 genes, we fit all five models using a hierarchical framework that pools evolutionary rate parameters across genes while permitting gene-specific optima, and includes a mixture component to account for genes that evolve without detectable phylogenetic signal (see Methods). We used simulations to verify that our inference procedure accurately recovers evolutionary parameters under each model (Supp Figs. S1, S2, S3, S4, S5)

### Comparative biophysical data support a burst-size-constrained model of expression evolution

After fitting all five phylogenetic models (using maximum likelihood) to the average log burst sizes, log(*b*), and log decay rates, log(*γ*), inferred from the Jiao et al. [44] data, we compared AIC values (**Fig. 3**). We found strong (and nearly identical) support for the *b*-constrained model (ΔAIC = 0 from best supported model) and the full-model (ΔAIC = 3); we note that when all the maximum likelihood parameters are inserted into their respective models, the H-matrices between the full and *b*-constrained model are structurally similar; the fact that they both have large inferred parameters in the second row shows that they both support strong tracking by *ℓ*_*γ*_. This implies that there are strong constraints on the evolution of *ℓ*_*b*_ that are independent of its contributions to the mean spliced mRNA level. The *γ*-constrained model was the next most supported model (ΔAIC = 40) and then the ESD model without any selection on the individual components (ΔAIC = 191). The independent model was far worse than any of these (ΔAIC = 777). The poor support for the independent model provides overwhelming evidence that the system components are indeed co-evolving exactly as predicted by the theory of ESD.

**Fig 3:**
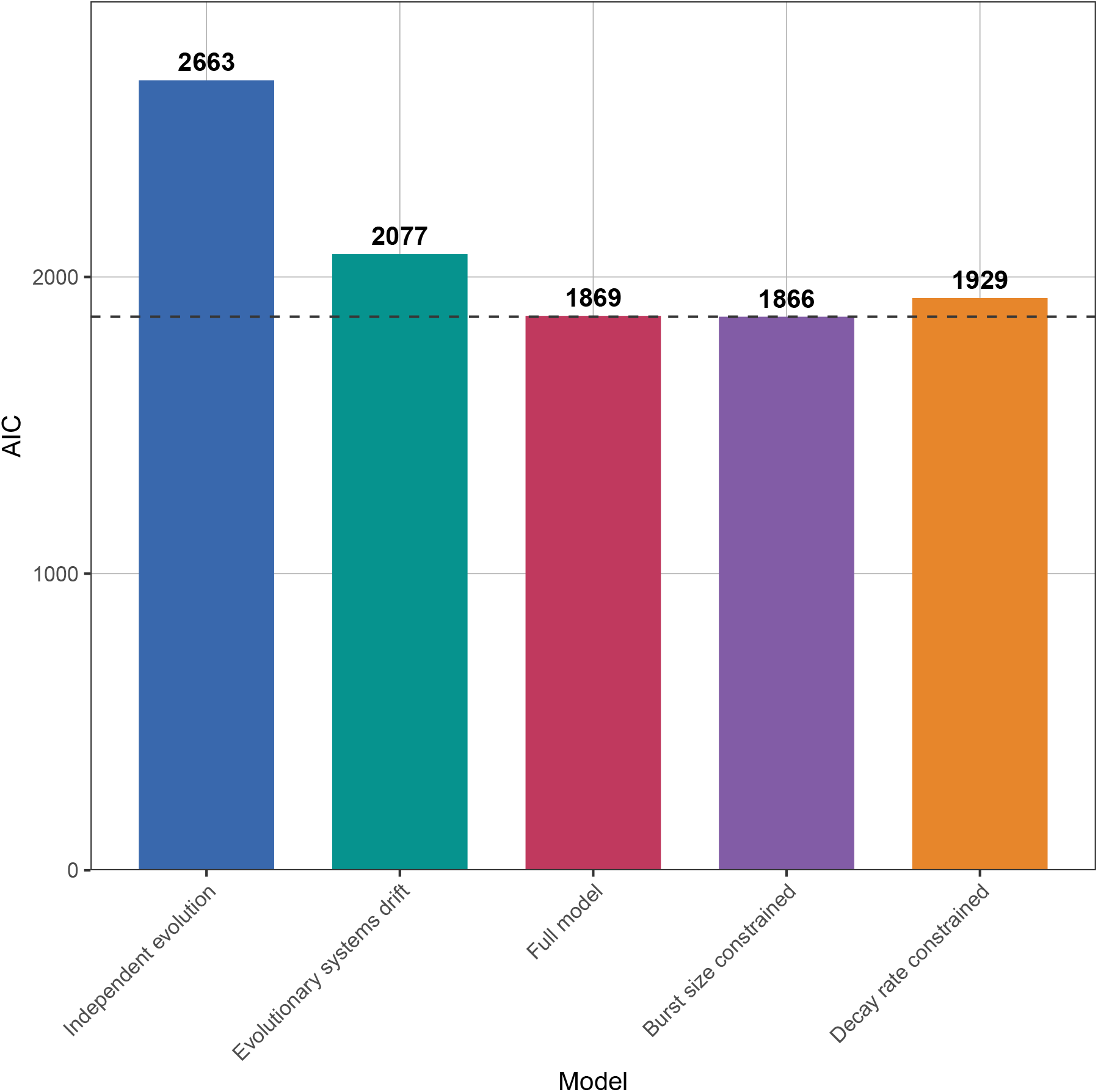
The burst-size constrained model best explains the co-evolution of biophysical parameters across vertebrates. AIC values for five competing phylogenetic models (lower AIC indicates better fit; dashed line marks the best-fitting model). The independent model, in which *b* and *γ* evolve without coupling, fits substantially worse than all models incorporating ESD-type coupling (ΔAIC = 586), indicating that the two parameters co-evolve in a coordinated manner as predicted by evolutionary systems drift. The pure ESD model, while considerably better than independent evolution, is itself outperformed by models that additionally include direct stabilizing selection on individual components (ΔAIC = 191 to the best model). The burst-size constrained model achieves the best fit, indicating that burst size faces strong direct stabilizing selection beyond what is required to maintain mean expression, while decay rate is primarily constrained through its contribution to mean expression

## Discussion

There are three primary contributions of this paper. First, we have estimated biophysical parameters of transcription and degradation across multiple species in a standardized way. To our knowledge, this has not been previously done. Then, by treating these parameters as quantitative traits that have evolved across the tree (see also, Neto-Bradley et al. [48], who applied a similar logic to physiological rates), we were able to gain richer insights into the processes driving divergence across species than we could by looking at mean spliced expression on its own. This approach is both general and scalable. Single-cell sequencing is becoming increasingly practical for non-model organisms [49–51] and it is clear that these datasets will become abundant in the near future. While we only considered a relatively simple biophysical model, the Monod framework [43] can accommodate a range of biophysical processes [30] and data modalities (such as ATAC-seq data [52] and proteomics [53]). Algorithmic advances in single-cell methods [54] mean that these models can be fit to the emerging large multi-species datasets. This opens up tremendous opportunities to understand how the biophysics of cellular processes have evolved.

Second, we have developed a novel quantitative formulation of ESD and discovered a signal for this process having played out across species. As cited above, many researchers have observed compensatory change between system components or simply component turnover with little effect on the phenotype; these findings have been at least partially attributable to the phenomenon of ESD. (Throughout this paper, we have taken an ahistorical perspective on ESD; tracing the history of the idea is complicated because it was developed several times apparently independently [e.g., [31, 32]], has been referred to by various names over the years, and many phenomena have only been retrospectively understood as falling under the umbrella of ESD.) Recent theoretical work [10, 41] revealed that for quantitative traits under stabilizing selection where the number of components is fixed, system components are expected to evolve marginally as a BM process. Empirically, Cope et al. [9] found that selective coupling between the mean protein level and mean mRNA level for a given gene was the most plausible explanation for why mean mRNA levels across species are inconsistent with Brownian evolution [11, 55]. Our new theoretical result is that, at least for the relatively simple cases we have studied where stabilizing selection acts directly on some linear combination of system components, the joint evolution of the components can be described via an OU process with a predictable correlation structure. As such, it is possible to fit such models to phylogenetic comparative data and evaluate the contribution of ESD per se to divergence among species. Our primary empirical result bears out the importance of ESD (Fig. 3). We find a very strong signal that the components of transcriptional systems have indeed drifted together in exactly the way we would predict under ESD and that this signal is independent of the signal for selective constraints on individual components.

More broadly, it is well-appreciated that, at least in finite populations, genetic drift is a background force that is shaping allele frequencies regardless of what else is happening in the population genetic environment. Our results suggest that when looking at the divergence of complex systems over macroevolutionary time (i.e., with sufficient time for compensatory mutations to occur and to fix [9]), researchers need to consider ESD as a persistent background effect against which inferences of adaptive processes are calibrated. Our results open up the possibility of doing so. There remain a number of open theoretical questions, such as how to model the evolution of systems with more complicated regulatory structures and how best to share information across observations (e.g., mRNA levels for different genes). It is also an open empirical question how important evolutionary systems drift is in shaping interspecific divergence across different systems and at different time scales; if there is tight mutational or selective coupling among components there will be limited systems drift [9]. Furthermore, as mutations accumulate in the system over long time scales, there are theoretical reasons that ESD may be expected to slow down [41]. We fully acknowledge that our species sampling was relatively small, that we were only able to accurately estimate parameter values for a limited number of genes, and that our conclusions are based on a single cell type (spleen T cells); given that this line of inquiry was only recently made possible by advances in single-cell biology (particularly in the advent of biophysical models [21]) and in evolutionary theory (including this work), we do not want to speculate as to how general these conclusions will be.

Third, our major empirical finding — in addition to the most direct evidence of ESD to date — is that macroevolutionary patterns of gene expression are consistent with strong selective constraints on transcriptional bursting per se that is independent of selection for mean spliced mRNA levels. We suggest that this is most plausibly due to selection to reduce transcriptional noise. This interpretation of the macroevolutionary data is consistent with experimental evidence within species. First, it is established (both in prokaryotes [56] and eukaryotes [57, 58]) that noise in mRNA expression is primarily affected by burst size and frequency [58, 59]; the mechanisms of mRNA decay, on the other hand, have a much less pronounced effect on the noisiness of expression [60, 61]. Note that in our biophysical model, we cannot distinguish between burst size and burst frequency, and hence what we instead consider is the combination of them as burst size. Second, there is evidence that the promoter sequences of “essential” or core genes show less noisy expression than other genes [62] and are subject to purifying selection [63]; genes that are more noisy tend to be those that are less conserved and those that are responsive to environmental conditions [64–66], suggesting a trade-off between noise reduction and evolutionary lability. This interpretation is further strengthened by recent evidence from both within- and between-species data that the variance of expression itself has a genetic variance (and is thus subject to selection) [67]. Our work suggests that these mechanisms scale up to generate large-scale patterns across vertebrates.

Overall, this work presents a new paradigm for investigating the evolutionary and genetic basis of interspecific divergence in gene regulation. Rather than looking only at mean mRNA levels (whether from bulk or single-cell data), our results demonstrate that researchers can investigate evolutionary changes in the biophysical parameters that describe various cellular processes. And with the ability to quantify the contribution of ESD to such changes, we can now disentangle the importance of adaptive from non-adaptive processes in shaping cellular systems across the tree of life.

## Methods

### Data processing and biophysical modeling

For this study, we used single-cell RNA-seq data from Jiao et al. [44], extracted from the spleen of seven vertebrate species. We processed the data using kallisto [68–71] to obtain spliced and unspliced count matrices. After clustering the data from each species by cell type, we excluded the fish sample from further analysis because of an indistinct and low-count T-cell cluster, leaving six vertebrate species for the spleen T-cell analysis. We searched for genes which had orthologs in all six retained species using Ensembl BioMart [72].

We then fit transcriptional rates for these genes in each species separately using Monod [43], using the bursty transcription model with Poisson technical noise. Monod is an inference framework which fits per-gene biophysical parameters for a selection of transcriptional models. The dynamics of the model are encapsulated in a chemical master equation, which can then be numerically solved to give steady-state distributions for spliced and unspliced RNA counts, including the impact of technical noise. The likelihood of the observed spliced/unspliced count matrices can then be maximized over biophysical parameters. In practice, this process is repeated over a grid of technical parameters, and the combined values which maximize the likelihood of the data are outputted.

The output of this procedure is a per-gene burst size, *b*, splicing rate *β*, and decay rate, *γ*, with the rates given in units of the transcription initiation rate, *k*. We then transformed these variables to log space. For simplicity in what follows, write *ℓ*_*b*_ = log(*b*), *ℓ*_*β*_ = log(*β*), and *ℓ*_*γ*_ = log(*γ*). We then subtracted the mean of each log-space parameter across genes from each species. After fitting with Monod, which filters some genes, and removing genes without a fitted ortholog in all six retained species, we were left with 167 genes. The mean-centered values of *ℓ*_*b*_, *ℓ*_*β*_ and *ℓ*_*γ*_ for each of these genes were used as the traits for the following analysis.

### Evolutionary dynamics under ESD

Previous work by Veller and Muralidhar [41] and Jiang et al. [10] showed that when selection acts only on the composite trait (i.e. the mean expression *µ*) rather than directly on the component traits (i.e., burst size *b* or degradation rate *γ*), then the component traits each evolve marginally as a BM, with no net directional pull on either parameter individually. In the Supplementary Material, we show that this nonetheless induces a correlation between the two components: intuitively, if the burst rate increases, then the degradation rate must also increase to maintain the same mean expression.

Thus, we were motivated to construct a bivariate model of systems drift to test if it could be distinguished from other modes of coevolution. Following the classical quantitative genetics framework of Lande [47], we assume that gene expression evolution can be described by stabilizing selection acting simultaneously on *ℓ*_*b*_, *ℓ*_*γ*_, and/or *ℓ*_*µ*_. Importantly, *ℓ*_*µ*_ = *ℓ*_*b*_ − *ℓ*_*γ*_. First, we consider a model where all selection acts through selection on *ℓ*_*µ*_

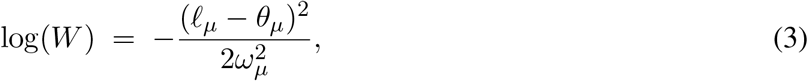

where *θ*_*µ*_ is the optimal level of gene expression and 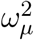 is the width of the fitness peak (*ω*_*µ*_ → ∞ corresponds to no selection on that quantity).

Let the genetic variance of *ℓ*_*µ*_ be *τ* ^2^, and denote by *f*_*b*_ the contribution to the genetic variance of *ℓ*_*µ*_ due to the genetic variance of *ℓ*_*b*_ and 1 − *f*_*b*_ the contribution from the genetic variance of *ℓ*_*γ*_. Then, the genetic covariance matrix of the vector **X**_*t*_ = (*ℓ*_*b*_, *ℓ*_*γ*_) is *G* = *τ* ^2^ diag(*f*_*b*_, 1 − *f*_*b*_). Then, we can use Lande’s formula [47] to see that **X**_*t*_ evolves as an OU process,

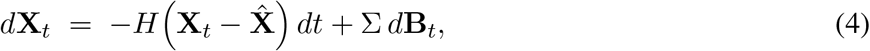

where *H* is the selection matrix and 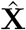 is the evolutionary equilibrium determined from the fitness function by 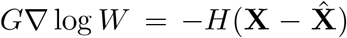. Σ is the cholesky square root of *G/N*, where *N* is the population size; that is, ΣΣ^⊤^ = *G/N*. For notational convenience, we define the compound parameter *σ*^2^ ≡ *τ* ^2^*/N* as the strength of genetic drift.

Under evolutionary systems drift, we find that the key compound parameter is

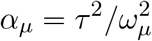

which measures the rate of adaptation of mean expression. Then, we can write the selection matrix

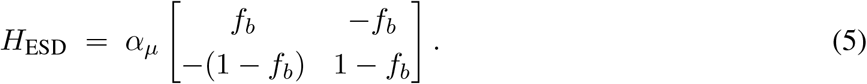

The insight is revealed by the eigendecomposition of *H*_ESD_:

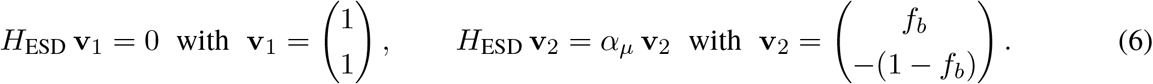

The zero eigenvalue corresponds to the direction **v**_1_ = (1, 1)^⊤^, along which *ℓ*_*b*_ and *ℓ*_*γ*_ change in exact tandem so that *ℓ*_*µ*_ = *ℓ*_*b*_ − *ℓ*_*γ*_ remains constant and incurs no fitness cost. Evolution along this neutral axis is a pure BM as noted by Veller and Muralidhar [41] and Jiang et al. [10]. The non-zero eigenvalue *α*_*µ*_ corresponds to the direction **v**_2_ ∝ (*f*_*b*_, −(1 − *f*_*b*_))^⊤^, along which *ℓ*_*µ*_ changes and is resisted by stabilizing selection. Along this constrained axis, *ℓ*_*µ*_ itself evolves as a scalar OU process:

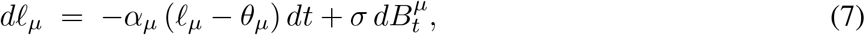

maintaining mean expression near its optimum *θ*_*µ*_ with selection strength *α*_*µ*_. The joint OU process therefore simultaneously captures two modes of evolution: free correlated drift of *ℓ*_*b*_ and *ℓ*_*γ*_ along **v**_1_, and OU-governed constraint on *ℓ*_*µ*_ along **v**_2_. Because *H*_ESD_ is singular (zero eigenvalue), the joint process has no stationary variance along **v**_1_. For calculations we regularize it by adding a small diagonal term *εI* (*ε* = 0.01).

### Five alternative models of evolution

The ESD model is one of five evolutionary scenarios we consider, each corresponding to different assumptions about where selection acts. To derive each model, we expand our fitness function (3) to the more general model

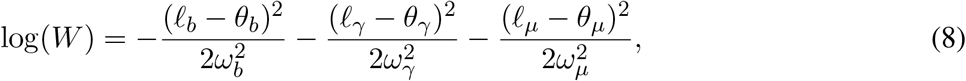

where *θ*_*i*_ is the optimum value of trait *ℓ*_*i*_ and 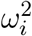 is the fitness width for trait *ℓ*_*i*_. We find, applying the same logic as before, we need to define two additional compound parameters,

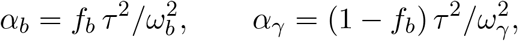

which denote the strength of direct stabilizing selection on *ℓ*_*b*_ and *ℓ*_*γ*_ respectively.

Thus, there are five parameters total in the richest model: *α*_*µ*_, *α*_*b*_, and *α*_*γ*_ measure the strength of selection on mean expression, burst rate, and degradation rate, respectively, while *σ*^2^ quantifies the rate of genetic drift and *f*_*b*_ is the proportion of the genetic variance of mean expression explained by the genetic variance of burst rate. Not all parameters are included in all models (see Table 1). The selection matrices for the five models are:

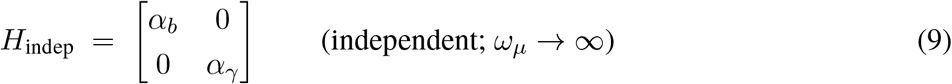

**Table 1:** Summary table of phylogenetic model fitted parameters: selection strengths, *α*_*b,γ,µ*_, contribution of *ℓ*_*b*_ to the genetic variance of *ℓ*_*µ*_ denoted by *f*_*b*_, stochastic parameters, *σ* (drawn separately for the independent model), along with the probability for a gene’s biophysical parameters to be drawn from a white-noise distribution, *p*_wn_ and the AIC of the model. Note that models have different parameters estimated based on the selective constraint hypothesis used for each model

| Model (Params) | $\alpha_b$ | $\alpha_\gamma$ | $\alpha_\mu$ | $f_b$ | $\sigma$ | $p_{wn}$ |
| --- | --- | --- | --- | --- | --- | --- |
| Full model (15) | 1.711 | 0.002 | 451.588 | 0.0247 | 5.368 | 0.2305 |
| Independent (14) | 2.269 | 3.989 | - | - | 0.696, 1.016 | 0.349 |
| ESD (9) | - | - | 421.577 | 0.0154 | 5.469 | 0.168 |
| γ-constrained (14) | - | 3.143 | 88.987 | 0.357 | 1.486 | 0.219 |
| $b$-constrained (14) | 1.825 | - | 588.937 | 0.0187 | 6.125 | 0.2002 |

The ESD model (*ω*_*b*_, *ω*_*γ*_ → ∞) is given by Equation (5), derived in full above.

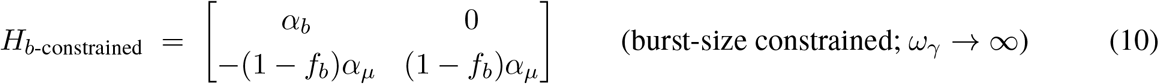

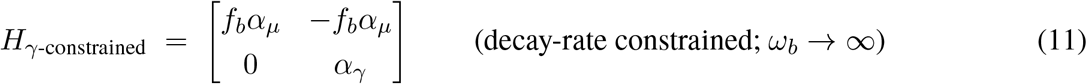

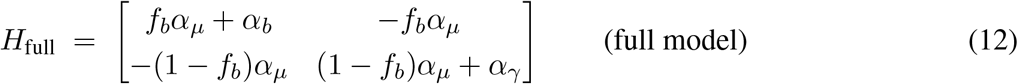

In the independent model (Eq. (9)), *ℓ*_*b*_ and *ℓ*_*γ*_ evolve toward their own optima without any interaction; there is no coupling induced by *ℓ*_*µ*_, and the off-diagonal entries are zero. The *b*- and *γ*-constrained models (Eqs. (10)–(11)) build on the ESD structure by adding direct stabilizing selection on one component: for example, in the *b*-constrained model, burst size is directly constrained (via *α*_*b*_) while decay rate is only indirectly constrained through its contribution to *µ* (via (1 − *f*_*b*_)*α*_*µ*_). The full model (Eq. (12)) combines ESD-type coupling with direct selection on both components.

All models except the independent model share the same diffusion matrix, *G/N* = *σ*^2^ diag(*f*_*b*_, 1−*f*_*b*_); in the independent model, 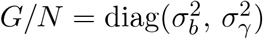, indicating that the two parameters drift independently with separate diffusion parameters.

### Parameter inference from phylogenetic comparative data

Likelihood equations for OU processes on phylogenetic trees are well-known [46, 73] (for the sake of completeness, we provide full-derivations of these likelihoods in the Supplementary materials). We used the PCMBase package [74] to implement the five variants we derived above. Because we have biophysical parameter estimates from 167 genes but only six species, we pool information hierarchically: all genes are assumed to share the same evolutionary rate parameters (*H, M*) but have gene-specific optima 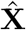, which are treated as random effects drawn from a multivariate Gaussian distribution. We integrate analytically over these gene-specific optima, yielding a tractable marginal likelihood over the evolutionary parameters. To account for genes that lack phylogenetic signal — arising from technical noise or rapid lineage-specific shifts — we include a mixture component: with probability *p*_wn_, a gene’s biophysical parameters are drawn from a white-noise (species-independent) distribution, and with probability 1 − *p*_wn_ from the OU model (following [75]). We used simulations to assess the suitability of our model for inferring evolutionary parameters from biophysical parameters. In all scenarios, we find that we accurately estimate parameters (Supp Fig. S1, S2, S3, S4, S5) All parameters are estimated by maximum likelihood [76, 77]; we compared models using the Akaike Information Criterion (AIC), with parameter estimates provided in Supp Mat S8.

## Data and code availability

The Genotype-Tissue Expression (GTEx) Project was supported by the Common Fund of the Office of the Director of the National Institutes of Health, and by NCI, NHGRI, NHLBI, NIDA, NIMH, and NINDS. The average expression data used for the gene binning described in this manuscript were obtained from the GTEx Portal, V10 spleen tissue, on 10/15/2025. The spleen single-cell transcriptomic data used in our analysis are taken from Jiao et al.[44]; after excluding the fish sample, our phylogenetic analysis used spleen T cells from six retained vertebrate species. Fig.2 - Created in BioRender. Krish, M. (2026) https://BioRender.com/ceagst3. The code and data for the phylogenetic model used to perform these analyses, and scripts to reproduce Fig 1, Fig. 3 and the supplementary figures are available at https://github.com/applied-phylo-lab/biophysics_esd_paper/tree/main [78].

## Acknowledgements

M.K. and M.P. were supported by NIGMS award R35GM151348 and startup funds from Cornell University. We also thank Charles Trimble for generously funding part of C.F.’s research through Caltech’s CI2 grant program.

## Author Contributions

Conceptualization, C.F., L.P., and M.P.; mathematical theory, C.F., J.G.S., M.K., and M.P.; computational analyses, C.F., M.K. (with guidance from J.G.S., A.L.C., L.P., and M.P.); writing–original draft, C.F.; writing– review and editing, C.F., J.G.S, M.K., A.L.C., L.P., and M.P.; funding acquisition–C.F. and M.P.; supervision– L.P. and M.P.

## Supplementary materials

### S1 Data processing

#### S1.1 Pre-processing

For this study, we used single-cell RNA-seq data from Jiao et al. [1], extracted from the spleen of seven vertebrate species. We processed the data using kallisto [2] to obtain spliced and unspliced count matrices, and filtered out low UMI cells. An example kallisto call is given here:

~~~
kb count --verbose -i ./frog/index.idx -g ./frog/t2g_mm10.txt -x 10xv2
    -o ./frog/output -t 24 -m 8G -c1 ./frog/cdna_t2c.txt -c2 ./frog/intron_t2c.txt
    --workflow=nac --filter bustools --strand=unstranded --sum=cell
    ../SRR16490736_1.fastq ../SRR16490736_2.fastq,
~~~

with example output summary:

~~~
“n_targets”: 76913,
“n_bootstraps”: 0,
“n_processed”: 289062079,
“n_pseudoaligned”: 190915538,
“n_unique”: 35185121,
“p_pseudoaligned”: 66.0,
“p_unique”: 12.2,
“kallisto_version”: “0.50.1”,
“index_version”: 13,
“start_time”: “Wed Jun 12 15:04:30 2024”
~~~

After clustering the data from each species by cell type, we excluded the fish sample from further analysis because of an indistinct and low-count T-cell cluster, leaving six vertebrate species for the spleen T-cell analysis.

#### S1.2 Fitting biophysical parameters

We searched for genes which had orthologs in all six retained species using Ensembl BioMart[3]. We then fit transcriptional rates for these genes in each species separately using Monod [4], using the bursty transcription model with Poisson technical noise. After fitting with Monod, which filters some genes, and removing genes without a fitted ortholog in all six retained species, we were left with 167 genes. An example of the Monod run is given below:

~~~
fitmodel = cme_toolbox.CMEModel(‘Bursty’,’Poisson’)
filt_param = {‘min_means’:[0.01, 0.01], ‘max_maxes’:[350, 350], ‘min_maxes’
;:[1,3]}
lb = [-1.0, -1.8, -1.8]
ub = [4.2, 2.5, 3.5]
# samp_lb, samp_ub = [-8, -3],[-5, 0]
samp_lb, samp_ub = [-11, -6],[-5, 0]
grid = [6,7]
fitted_adata = inference.perform_inference(combined_adata, fitmodel, n_genes=5000, seed=5, phys_lb=lb, phys_ub=ub, gridsize=grid, samp_lb=samp_lb, samp_ub=samp_ub, filt_param=filt_param, gradient_param={‘max_iterations’:5,’init_pattern’:’moments’,’num_restarts’:1}, dataset_string=dataset_string, viz=True,num_cores=32)
~~~

The output of this procedure is a per-gene burst size, *b*, splicing rate *β*, and decay rate, *γ*, with the rates given in units of the transcription initiation rate, *k*, all in log space. We then subtracted the mean of each parameter across genes from each species, and used the resulting values as the traits for the phylogenetic analysis.

#### S1.3 Phylogenetic tree

For the phylogenetic tree, we used the following tree, from TimeTree [5], in Newick format:

~~~
(Frog:351.68654000,(Pig:94.00000000,((Rat:11.64917000,Mouse:11.64917000) ‘14’:75.55083000,(Human:28.82000000,Macaque:28.82000000)’13’:58.38000000) ‘25’:6.80000000)’37’:257.68654000);
~~~

### S2 Evolutionary systems drift

The quantitative systems drift theory from Veller and Muralidhar [6] shows that any component of a trait can drift marginally as if unconstrained. Here, we show that systems drift will induce a correlation between a pair of components, indicating that it may be possible to distinguish systems drift from pure drift by looking at multivariate evolutionary models.

Note that, since the unspliced mean, *µ*_*u*_, is given by 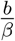, and the spliced mean is given by 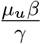, the splicing rate *β* does not influenced the mean spliced counts, *µ*_*s*_ (recall that rates are fit in units of the transcriptional initiation rate). Hence we can write,

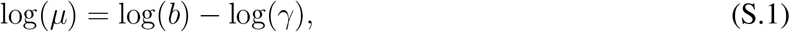

For notational simplicity, in the rest of this supplementary material we will write *ℓ*_*b*_ = log(*b*), *ℓ*_*γ*_ = log(*γ*), and *ℓ*_*µ*_ = log(*µ*).

We begin with the composite trait *ℓ*_*µ*_ = *ℓ*_*b*_ − *ℓ*_*γ*_, which we assume satisfies an Ornstein-Uhlenbeck (OU) process according to:

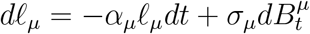

Following Veller and Muralidhar, we split this up into two component equations according to Equation (S.1) and that *ℓ*_*b*_ accounts for a proportion *f*_*b*_ of the genetic variance for *ℓ*_*µ*_ while *ℓ*_*γ*_ accounts for a proportion 1 − *f*_*b*_ of the genetic variance:

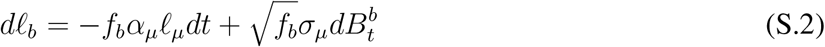

and

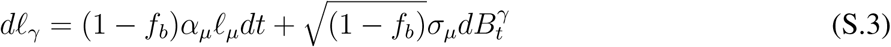

where it can be verified that *dℓ*_*µ*_ = *dℓ*_*b*_ − *dℓ*_*γ*_, where 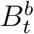 and 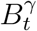 are standard Brownian motions such that 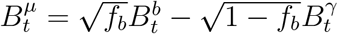

Using the same strategies found in their paper [6], we transform Equations (S.2) and (S.3) into

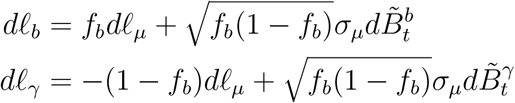

where

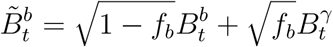

and

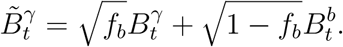

The integrated forms then become

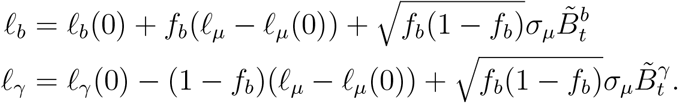

Notably, we see that 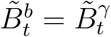 so that

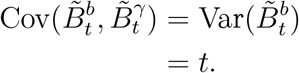

Thus, using independence of 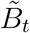 from *ℓ*_*µ*_ [6] and our previous result about the covariance of the Brownian motions,

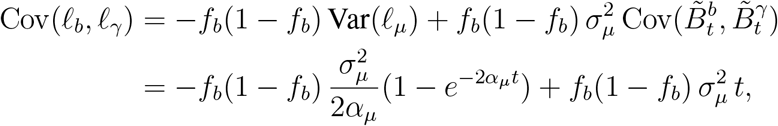

where we have also used the standard result for the variance of an OU process. For large *t*, the OU variance term saturates and the covariance is dominated by the linearly growing term:

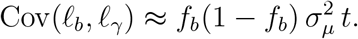

Thus, systems drift induces a dependence between the two components: even if marginally they appear to drift freely, by examining them *jointly* we can determine whether they are drifting independently or whether they are constrained by their contributions to *ℓ*_*µ*_.

### S3 Deriving the evolutionary dynamics as an Ornstein Uhlenbeck equation

Motivated by the previous section showing that systems drift induces a correlation between the drifting components, we now derive a joint model of systems drift.

For maximum generality, we start with the assumption that selection acts on the component traits (*ℓ*_*b*_ and *ℓ*_*γ*_) as well as the overall trait, in this case the mean spliced mRNA level *ℓ*_*µ*_. With this, we consider the following fitness function:

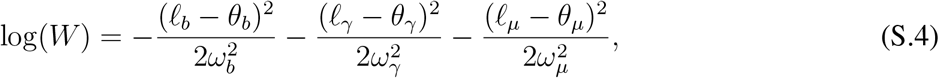

where

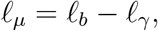

*θ*_*i*_ represents the evolutionary optimum of trait *ℓ*_*i*_, and 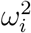 is the selective width for trait *i*. If 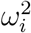 is larger, then selection is weaker.

Using classical evolutionary quantitative genetics theory [7], the bivariate process **X**_*t*_ = (*ℓ*_*b*_, *ℓ*_*γ*_)^⊤^, evolves according to

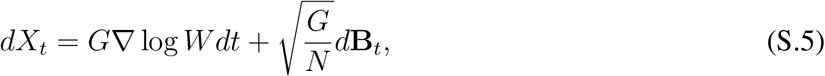

where *G* is the genetic covariance matrix. Note that 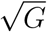 is the Cholesky square root of *G*. We take

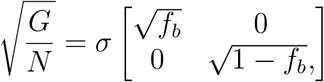

indicating that *ℓ*_*b*_ accounts for a proportion *f*_*b*_ of the genetic variance for *ℓ*_*µ*_ while *ℓ*_*γ*_ accounts for a proportion 1 − *f*_*b*_ of the genetic variance. Then, for convenience, we define 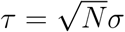 so that

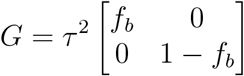

Comparing Equation (S.5) to the standard multivariate Ornstein Uhlenbeck form [8, 9]:

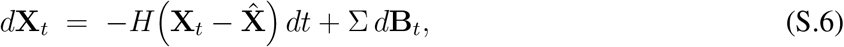

we determine *H*, 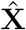 and Σ by equating

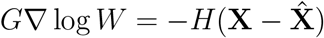

to find that

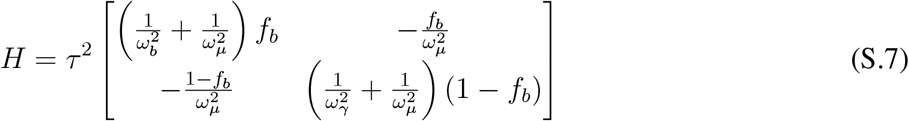

and

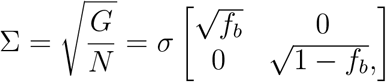

so that the diffusion matrix is

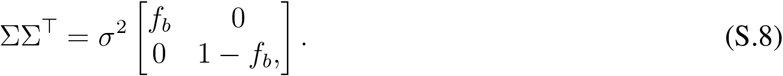

Next, define the compound parameters

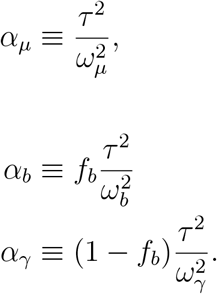

Then, using these compound parameters, we recover the **full model**,

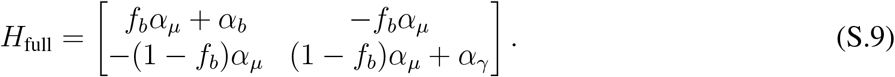

#### S3.1 Submodels

We now consider all the submodels of the full model that are discussed in the main text.

First, by turning off selection on each component trait individually, i.e. sending 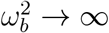 and 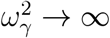 (or equivalently sending *α*_*b*_ → 0 and *α*_*γ*_ → 0), we recover the **systems drift model**,

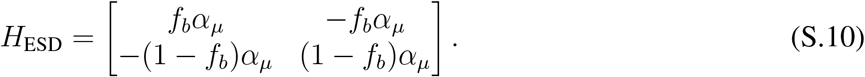

Similarly, by turning off selection on the composite trait *ℓ*_*µ*_, by sending 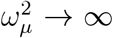 (equivalently *α*_*µ*_ → 0), we obtain the **independent model**,

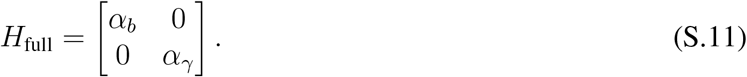

To obtain the two constrained models, we need to take slightly more complex limits, because we need selection on one component to dominate the dynamics, while the other component is selected only to maintain mean expression. To obtain the b-constrained model, rewrite Equation (S.9) as

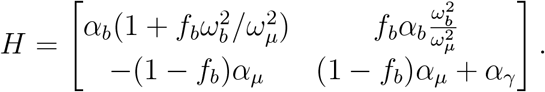

Then, we take *α*_*γ*_ → 0 to turn off direct selection on *ℓ*_*γ*_ and send 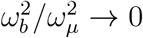 to indicate that selection on *ℓ*_*b*_ is significantly stronger than selection on *ℓ*_*µ*_. This yields the *b***-constrained** model,

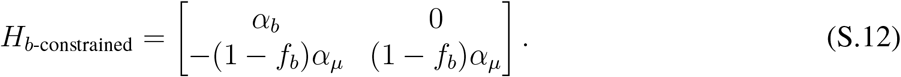

The same approach, mutatis mutandis, will achieve the *γ***-constrained** model,

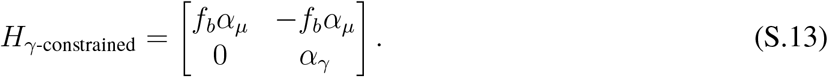

### S4 Multi-OU on a tree with distribution on the root state

Our approach pools information across genes in order to gain power from a small tree [10]. However, while the evolutionary parameters may be similar across genes, it is unlikely that the root and equilibrium traits are the same across genes. Hence, we incorporate a distribution over root states and equilibrium values to capture variation among genes [11]. Here, we briefly recall some central results and outline our approach.

First, recall S.6:

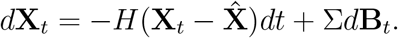

It can be shown [8, 9] that the solution is

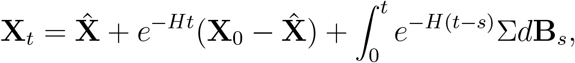

which is Gaussian with mean

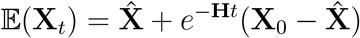

and variance

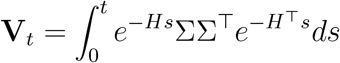

which also satisfies the differential equation

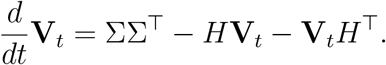

If a stationary distribution exists, the stationary variance satisfies the Lyapunov equation

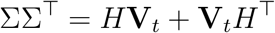

In practice, we make use of the identity

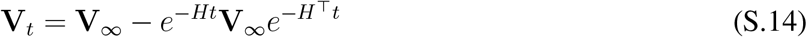

to compute the variance of the process. This can be seen because

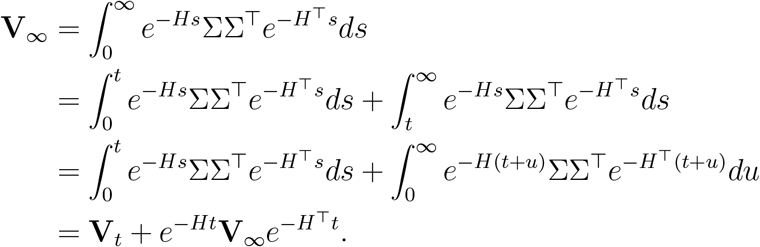

With a fixed equilibrium value 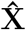 and root **X**_0_, the covariance between tips *i* and *j* is

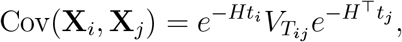

where *T*_*ij*_ is the root-to-MRCA time and *t*_*i*_, *t*_*j*_ are branch lengths from the MRCA to each tip. Further, because we assume the tree is ultrametric, *T* = *T*_*ij*_ + *t*_*i*_ is the height of the tree.

#### S4.1 Distribution on the root state

Suppose that the root state has mean *µ*_0_ and variance *V*_0_. Then, using the law of total expectation,

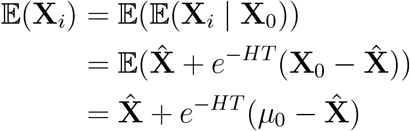

while the covariance given fixed 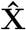 is

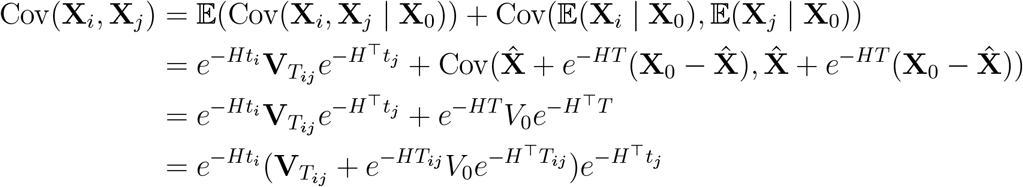

where we have used *T* = *T*_*ij*_ + *t*_*i*_.

We now also include a distribution of equilibrium values, 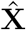. Given a value of 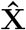, we assume that the distribution of the root state is centered on 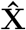. Hence, 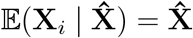 and by the law of total expectation

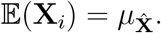

Then, by the law of total covariance,

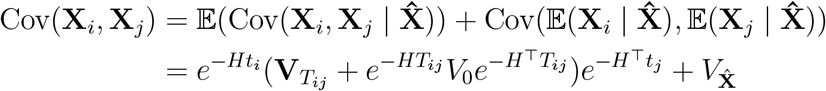

Using Equation (S.14) to compute this as

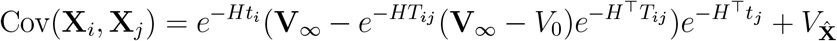

For simplicity, we assume the root variance is diagonal, that is

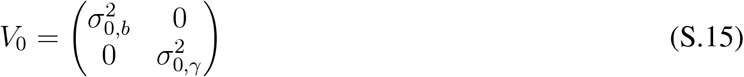

where *σ*_0,*b*_ and *σ*_0,*γ*_ refer to the standard deviation of *ℓ*_*b*_ and *ℓ*_*γ*_ respectively at the root.

#### S4.2 Covariance of the equilibrium distribution

For all models besides systems drift, we simply assume that the distribution of equilibrium values is Gaussian with mean

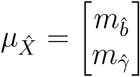

and covariance matrix

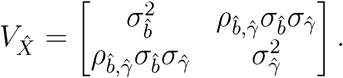

The systems drift model is more highly constrained, because there is only one optimum, which is *θ*_*µ*_. This forces a self-consistent quasi-optimum distribution on the components *ℓ*_*b*_ and *ℓ*_*γ*_, which we compute here. First, note that under the systems drift model

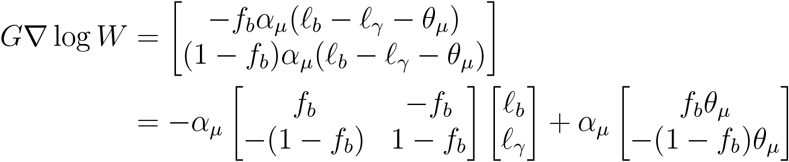

To write this in the form 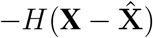 we must have

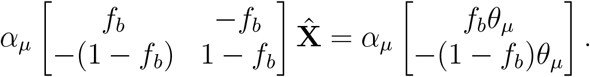

This is underdetermined, and any choice such that

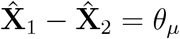

will work. A simple choice is

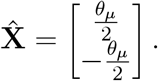

Thus, if we have that *θ*_*µ*_ is Gaussian distributed with mean 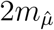 and variance 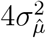, we have that the **systems drift model** can be parameterized as a degenerate 2-dimensional Gaussian with mean

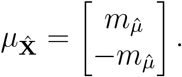

and covariance matrix

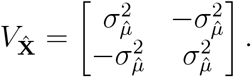

Again, we note that this is not tantamount to assuming a stationary distribution at the root, instead this simply provides a self-consistent prior distribution on the quasi-optimum values for the components that is consistent with a distribution on the optimum of the composite.

### S5 Computational considerations

#### S5.1 Parameterizing the optimization geometry

Although the parameterizations we described above are informative about the dynamics of the system, they are somewhat difficult to optimize due to the appearance of terms like *f*_*b*_*α*_*µ*_. Hence, for optimization, we instead define new compound parameters

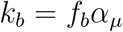

and

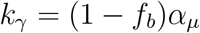

and use these as our optimization parameters in the *H* matrix. Note that under this parameterization, we parameterize the diffusion matrix as

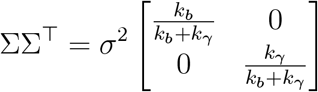

in order to maintain the correct ties between the parameters, except for the independent model, in which 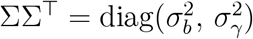.

When reporting our results in the main text, we convert back to the original parameterization, with *α*_*µ*_ and *f*_*b*_, because the transformation is one-to-one.

#### S5.2 Regularization of the Systems Drift model

Since the H given in Equation (S.10) is singular, we cannot naively use the formula (S.14) to compute that variance. Instead, we regularize via the addition of a small diagonal matrix as:

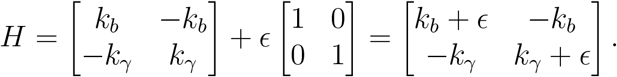

Solving for **V**_∞_ using this H matrix gives us:

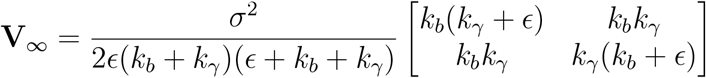

For further analysis, we fix the value of *ϵ* = 0.01. We note that this is purely done to aid in computation, and as *ϵ* → 0 it has no impact on the results, as we are not assuming that the process starts from the stationary distribution.

### S6 Simulation and data-fit results

For the model comparison, we simulate 100 datasets under each of the five hypotheses using PCMBase [12]. We draw random sets of the respective true parameters for each simulation uniformly between the bounds. We then fit the simulated datasets using the models derived. The simulation results and the parameter estimates for the data are shown respectively for each of the five models: Independent evolution (Supp Fig. S1, Supp Table 1), Evolutionary systems drift (Supp Fig. S2, Supp Table 2), Full model (Supp Fig. S3, Supp Table 3), Burst size constrained model (Supp Fig. S4, Supp Table 4), and Decay rate constrained model (Supp Fig. S5, Supp Table 5).

### Supplementary Figures and Tables

**Figure S1:**
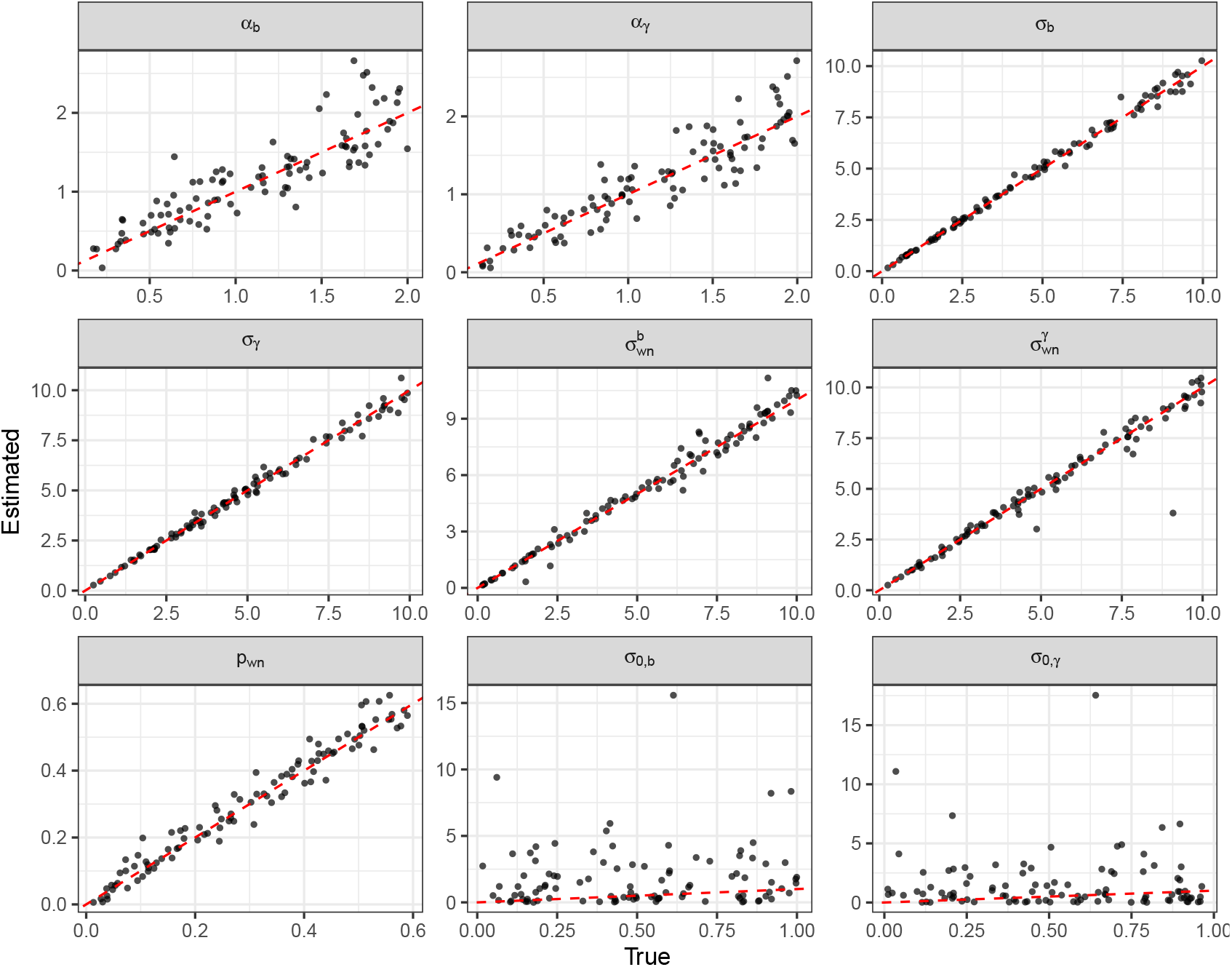
Fitted vs estimated parameters for independent evolution model

**Table 1:** Estimated parameters for the Independent model fitted to the data.

|  |  |  |  |  |  |  |  |
| --- | --- | --- | --- | --- | --- | --- | --- |
| $\alpha_b$ | $\alpha_\gamma$ | $\sigma_b$ | $\sigma_\gamma$ | $\sigma_{wn}^b$ | $\sigma_{wn}^\gamma$ | $p_{wn}$ | |
| 2.269 | 3.989 | 0.696 | 1.016 | 0.634 | 0.697 | 0.349 |  |
| $m_{\hat{b}}$ | $\sigma_{\hat{b}}$ | $m_{\hat{\gamma}}$ | $\sigma_{\hat{\gamma}}$ | $\rho_{\hat{b},\hat{\gamma}}$ | $\sigma_{0,b}$ | $\sigma_{0,\gamma}$ | AIC |
| 2.49e-04 | 0.218 | 0.025 | 0.619 | 0.994 | 0.549 | 0.263 | 2663 |

**Figure S2:**
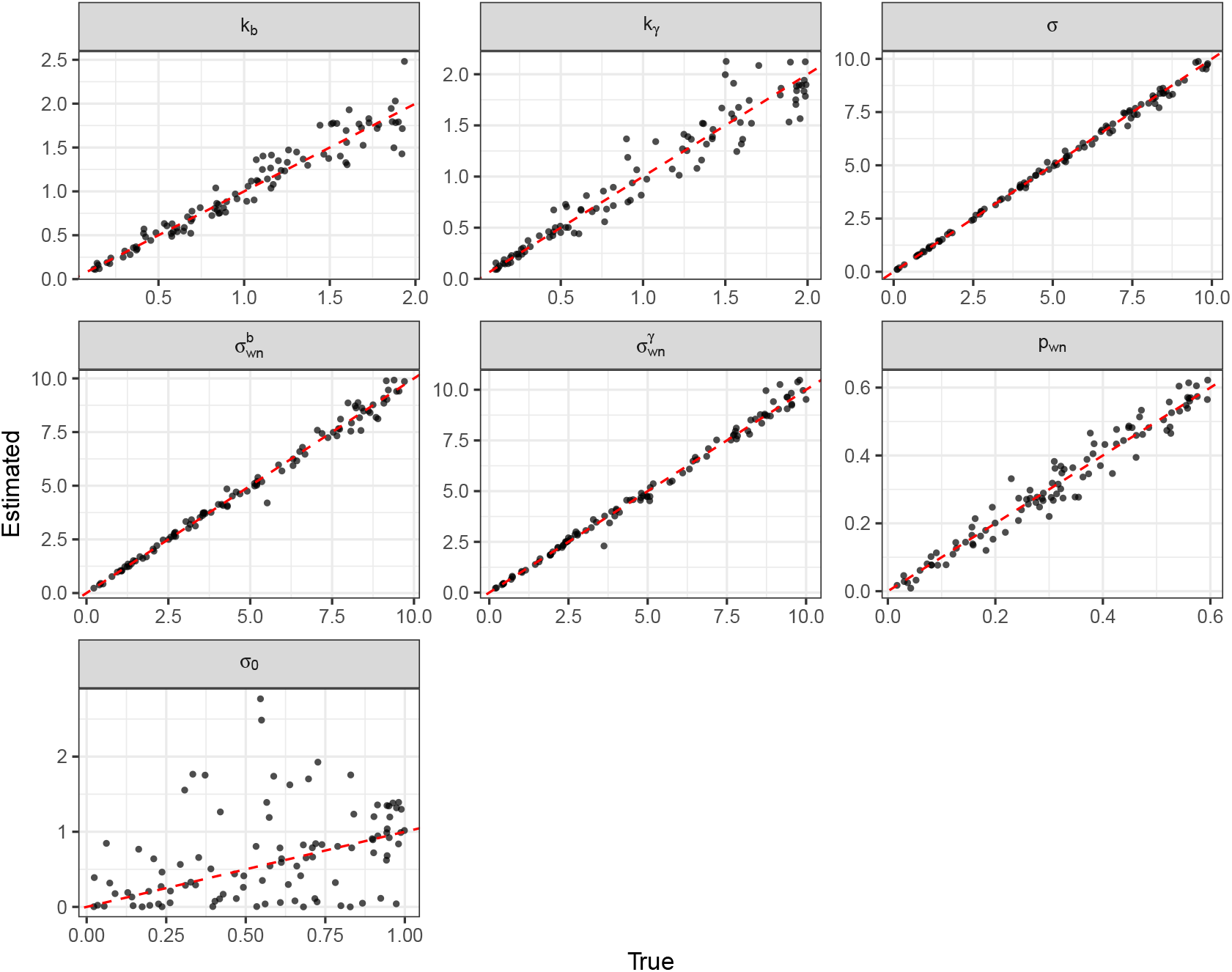
Fitted vs estimated parameters for Evolutionary systems drift model

**Table 2:** Estimated parameters for the Evolutionary Systems Drift model fitted to the data.

|  |  |  |  |  |  |  |  |  |  |
| --- | --- | --- | --- | --- | --- | --- | --- | --- | --- |
| $k_b$ | $k_\gamma$ | $\sigma$ | $\sigma_{wn}^b$ | $\sigma_{wn}^\gamma$ | $p_{wn}$ | $m_{\hat{\mu}}$ | $\sigma_{\hat{\mu}}$ | $\sigma_0$ | AIC |
| 6.488 | 415.089 | 5.469 | 0.709 | 1.269 | 0.168 | -0.101 | 0.181 | 2.6e-06 | 2077 |

**Table 3:** Estimated parameters for the Full model fitted to the data.

|  |  |  |  |  |  |  |  |  |
| --- | --- | --- | --- | --- | --- | --- | --- | --- |
| $\alpha_b$ | $\alpha_\gamma$ | $k_b$ | $k_\gamma$ | $\sigma$ | $\sigma_{wn}^b$ | $\sigma_{wn}^\gamma$ | $p_{wn}$ | |
| 1.7114 | 0.0019 | 11.199 | 440.388 | 5.368 | 0.6929 | 0.7464 | 0.2305 |  |
| $m_{\hat{b}}$ | $\sigma_{\hat{b}}$ | $m_{\hat{\gamma}}$ | $\sigma_{\hat{\gamma}}$ | $\rho_{\hat{b},\hat{\gamma}}$ | $\sigma_{0,b}$ | $\sigma_{0,\gamma}$ | | AIC |
| -0.0141 | 0.1665 | 0.0553 | 0.5773 | 0.957 | 0.7323 | 4.2e-05 |  | 1869 |

**Table 4:** Estimated parameters for the Burst size constrained model fitted to the data.

|  |  |  |  |  |  |  |  |
| --- | --- | --- | --- | --- | --- | --- | --- |
| $\alpha_b$ | $k_b$ | $k_\gamma$ | $\sigma$ | $\sigma_{wn}^b$ | $\sigma_{wn}^\gamma$ | $p_{wn}$ | |
| 1.825 | 11.033 | 577.904 | 6.125 | 0.686 | 0.783 | 0.2002 |  |
| $m_{\hat{b}}$ | $\sigma_{\hat{b}}$ | $m_{\hat{\gamma}}$ | $\sigma_{\hat{\gamma}}$ | $\rho_{\hat{b},\hat{\gamma}}$ | $\sigma_{0,b}$ | $\sigma_{0,\gamma}$ | AIC |
| -3.9e-03 | 0.243 | 0.053 | 0.596 | 0.857 | 0.067 | 0.115 | 1866 |

**Figure S3:**
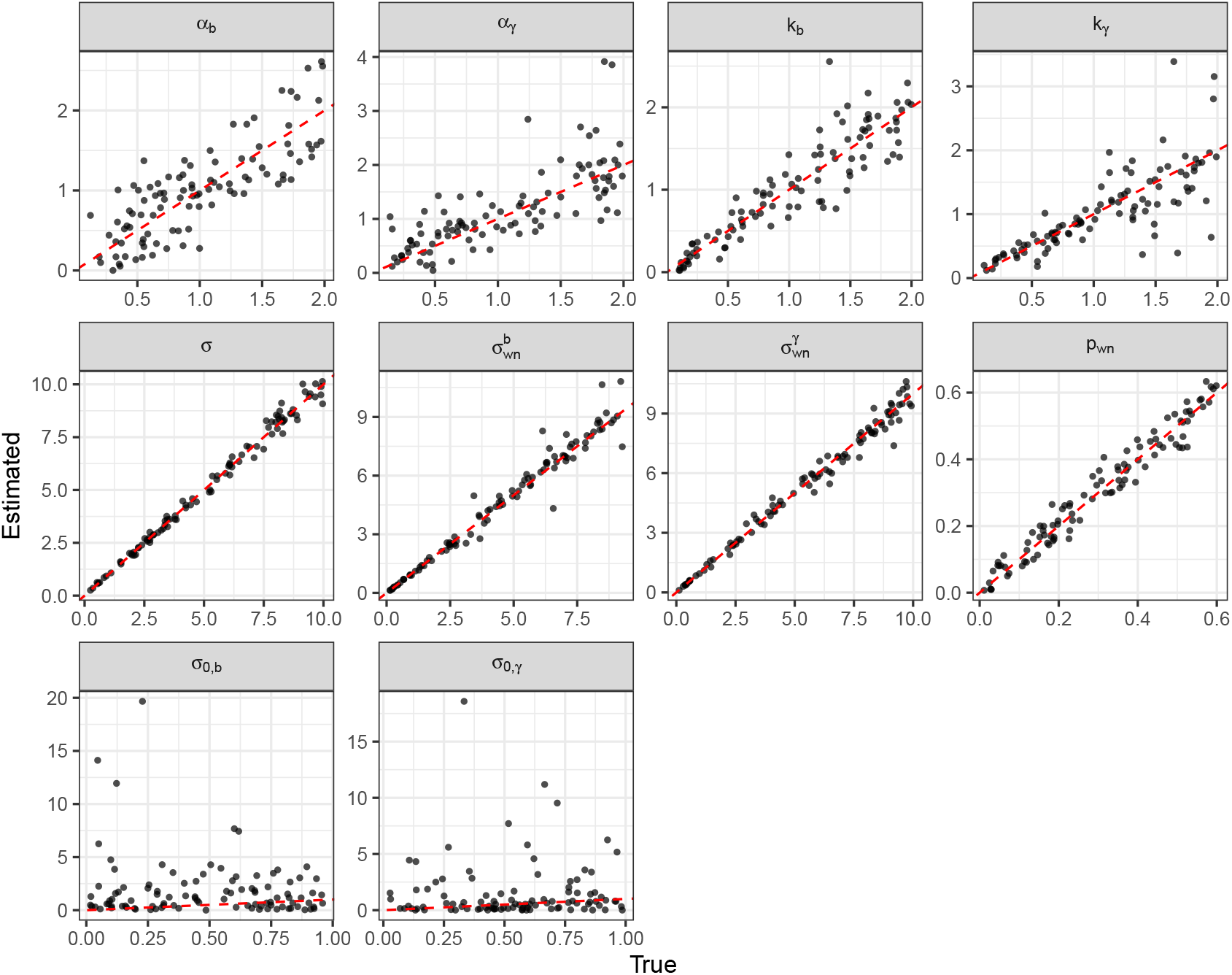
Fitted vs estimated parameters for the full model

**Figure S4:**
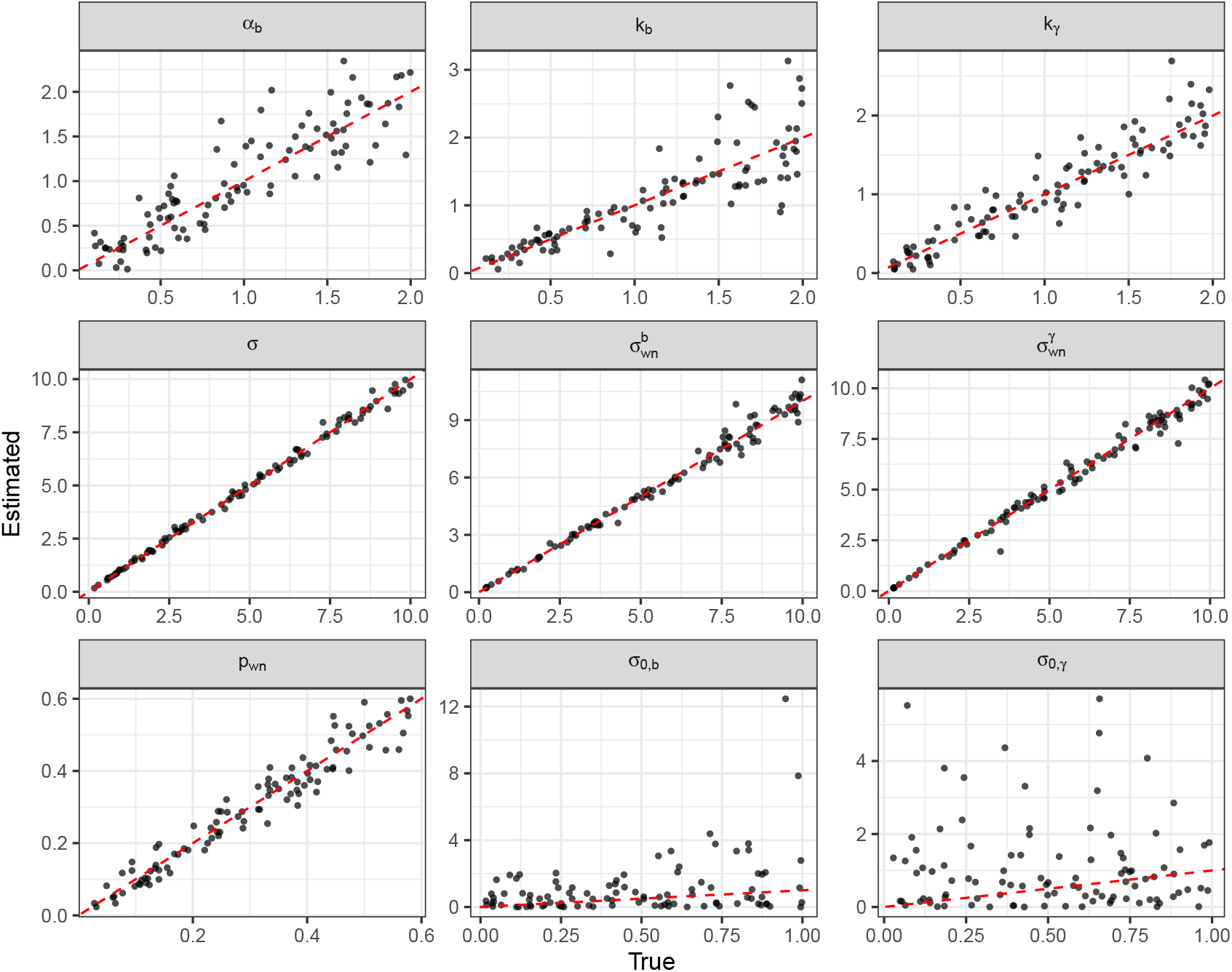
Fitted vs estimated parameters for burst size constrained model

**Figure S5:**
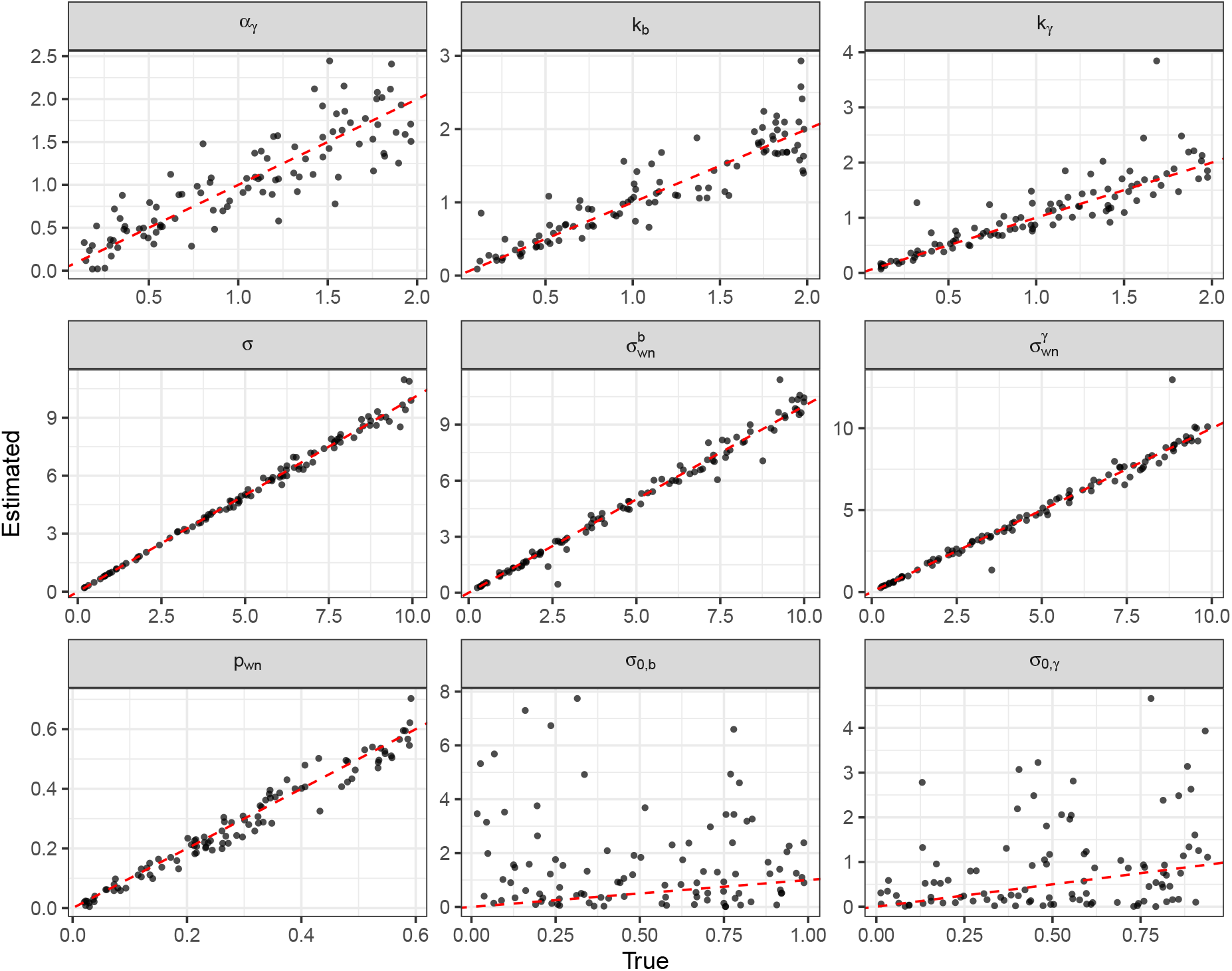
Fitted vs estimated parameters for decay rate constrained model

**Table 5:** Estimated parameters for the Decay rate constrained model fitted to the data.

| $\alpha_\gamma$ | $k_b$ | $k_\gamma$ | $\sigma$ | $\sigma_{wn}^b$ | $\sigma_{wn}^\gamma$ | $p_{wn}$ | |
| --- | --- | --- | --- | --- | --- | --- | --- |
| 3.143 | 31.789 | 57.198 | 1.486 | 0.670 | 0.780 | 0.219 |  |
| $m_{\hat{b}}$ | $\sigma_{\hat{b}}$ | $m_{\hat{\gamma}}$ | $\sigma_{\hat{\gamma}}$ | $\rho_{\hat{b},\hat{\gamma}}$ | $\sigma_{0,b}$ | $\sigma_{0,\gamma}$ | AIC |
| -0.021 | 0.195 | 0.037 | 0.572 | 0.904 | 0.006 | 0.047 | 1929 |

